# Allele-specific antisense oligonucleotide therapy for dominantly inherited hearing impairment DFNA9

**DOI:** 10.1101/2020.09.29.316364

**Authors:** Erik de Vrieze, Jolien Peijnenborg, Jorge Cañas Martin, Aniek Martens, Jaap Oostrik, Simone van der Heuvel, Kornelia Neveling, Ronald Pennings, Hannie Kremer, Erwin van Wijk

**Author notes:** Correspondence should be addressed to E.d.V. Experiments were conducted at the Radboud University Medical Center in Nijmegen, the Netherlands., *Corresponding authors address:* Radboud University Medical Center, Dept of Otorhinolaryngology and Dept of Human Genetics, P.O. box 9101, 6500 HB Nijmegen (route 855), E.

## Abstract

The c.151C>T founder mutation in *COCH* is a frequent cause of late onset, dominantly inherited hearing impairment and vestibular dysfunction (DFNA9) in the Dutch/Belgian population. The initial clinical symptoms only manifest between the 3rd and 5th decade of life, which leaves ample time for therapeutic intervention. The dominant inheritance pattern and established non-haploinsufficiency disease mechanism indicate that suppressing translation of mutant *COCH* transcripts has high therapeutic potential. Single-Molecule Real-Time (SMRT) sequencing resulted in the identification of 11 variants with a low population-frequency (< 10%), that are specific to the c.151C>T mutant *COCH* allele. Proof of concept was obtained that gapmer antisense oligonucleotides (AONs), directed against the c.151C>T mutation or mutant allele-specific intronic variants, are able to specifically induce mutant *COCH* transcript degradation when delivered to transgenic cells expressing *COCH* minigenes. Sequence optimization of the AONs against the c.151C>T mutation resulted in a lead molecule that reduced the levels of mutant *COCH* transcripts by ~60% in a transgenic cell model, without affecting wildtype *COCH* transcript levels. With the proven safety of AONs in humans, and rapid advancements in inner ear drug delivery, our in-vitro studies indicate that AONs offer a promising treatment modality for DFNA9.

## Introduction

DFNA9, caused by mutations in the *COCH* gene, is a relatively common form of dominantly inherited highly progressive hearing loss and vestibular dysfunction. It is characterized by adult-onset hearing loss, leading to complete deafness by the age of 50-70 years ^1,2^. With progression of the disease, speech perception and conversation become severely limited. DFNA9 patients furthermore suffer from balance problems, which severely hamper their daily activities. Overall, the problems associated with DFNA9 have a severe impact on the quality of life of patients, their relatives and friends ^3^.

The *COCH* gene is located on chromosome 14, and encodes cochlin, a protein that consists of 550 amino acids. Cochlin is predicted to contain a signal peptide, an LCCL (Limulus factor C, Cochlin, and late gestation lung protein Lgl1) domain, two short intervening domains, and two vWFA (von Willebrandfactor A) domains. Cochlin is expressed in fibrocytes of the spiral ligament and spiral limbus, where it has been reported to assist in structural support, sound processing, and in the vestibular fibrocytes where is important in the maintenance of balance ^4^. Proteolytic cleavage of cochlin, between the LCCL domain and the more C-terminal vWFA domains, results in a 16-kDa LCCL domain-containing peptide that is secreted and has been shown to play a role in innate immunity in the cochlea ^5^. The vWFA domain-containing cochlin fragments are presumed to be extracellular matrix proteins, as cochlin vWFA2 was found to interact with collagens in-vitro, and cochlin is a major component of the cochlear extracellular matrix ^1,6^.

The c.151C>T (p.Pro51Ser) founder mutation, affecting the LCCL domain, appears to be the most prevalent mutation in *COCH*, as it underlies hearing loss in >1000 Dutch and Belgian individuals ^7,8^. Histopathology of a temporal bone from a p.Pro51Ser DFNA9 patient revealed significant loss and degeneration of fibrocytes in the cochlea (Robertson et al., 2006). Overexpression of murine cochlin containing the orthologue of the p.Pro51Ser variant in cultured cells, previously revealed that this mutation results in the formation of cytotoxic cochlin dimers and oligomers that sequester wildtype cochlin ^9^. While the proteolytic cleavage of cochlin was shown to be reduced by the p.Pro51Ser variant, and abolished by several other DFNA9-associated variants ^9^, the potential contribution of decreased proteolytic cleavage to DFNA9 pathology requires further investigation.

All available data indicates that DFNA9 results from a gain-of-function and/or a dominant-negative disease mechanism, rather than from haploinsufficiency. Downregulation of the mutant allele, thereby alleviating the inner ear from the burden caused by the formation of cytotoxic cochlin dimers, therefore has high therapeutic potential. The lack of auditory and vestibular phenotypes in mice carrying a heterozygous protein-truncating mutation in *Coch* ^10^, and in heterozygous family members of patients with early-onset hearing impairment due to homozygous protein-truncating mutations in *COCH* ^11^, illustrate that sufficient functional cochlin proteins can be produced from a single healthy *COCH* allele. We speculate that a timely intervention might even prevent hearing impairment altogether.

Antisense oligonucleotides with DNA-like properties can be employed to target (pre-)mRNA molecules for degradation by the RNase H1 endonuclease ^12,13^. Chemical modifications can be introduced in the 5’ and 3’ flanking nucleotides to increase stability and nuclease resistance, whilst maintaining a central gap region of oligo-deoxynucleotides to bind to the target RNA and thereby activate RNase H1 ^12^. These AONs are named gapmers, and their ability to specifically target mutant alleles for degradation has shown great promise in treatment strategies for non-haploinsufficiency disorders such as Huntington’s disease ^14,15^.

For a successful application of AON therapy for non-haploinsufficiency disorders such as DFNA9 it is of major importance that the designed AONs only target the mutant (pre-)mRNA, and not the wildtype (pre-)mRNA, for degradation. As the options to design allele-discriminating AONs based on a single nucleotide difference are limited, we used Single-Molecule Real-Time (SMRT) sequencing to identify additional allele-discriminating variants that can be exploited for AON design. This resulted in the identification of 11 variants with a low population frequency (< 10%), that are specific to the c.151C>T mutant *COCH* haplotype. Our results show that both the c.151C>T mutation in *COCH,* and low-frequency variants in *cis* with the DFNA9 mutation, can be used to specifically target mutant *COCH* transcripts for degradation by RNase H1. Lead molecule c.151C>T AON-E appears to be the most promising molecule for further preclinical investigation. As this AON targets the DFNA9-causing mutation, future clinical application is not limited by the potential presence of the target on the patient’s wildtype allele.

## Results

### Identification of therapeutic targets

In order to develop a mutant allele-specific therapy for DFNA9, reliable discrimination between the mutant and the wildtype allele is of vital importance. However, the single nucleotide changes in *COCH* underlying most cases of DFNA9, restrict the design of allele-discriminating therapies. In search of additional variants that can be exploited to improve discrimination between the c.151C>T mutant and wildtype *COCH* allele, we subjected the genomic *COCH* sequence of three DFNA9 patients to long-read single-molecule real-time (SMRT) sequencing. We amplified the *COCH* gene in three fragments that contain overlapping SNPs (c.151C>T and c.734-304T>G) to aid haplotype assembly (Figure 1A). The identified variants are annotated on transcript NM_001135058.1, which does not contain the extended second coding exon. To identify targetable allele-specific variants that potentially allow for the treatment of the majority of the Dutch/Belgian DFNA9 patients, we filtered the variants *in cis* with the c.151C>T mutation for a population frequency below 20%. This resulted in the identification of 11 deep-intronic variants, that are specific for the c.151C>T mutant *COCH* allele, and have allele frequencies between 5% and 10% (Figure 1B; Table 1). The identified variants provide additional targets for the development of a mutant allele-specific genetic therapy. The identified variants were validated using Sanger sequencing, and confirmed to segregate with the c.151C<T mutation in *COCH* in two branches of Dutch DFNA9 families (Figure S1).

**Figure 1.**
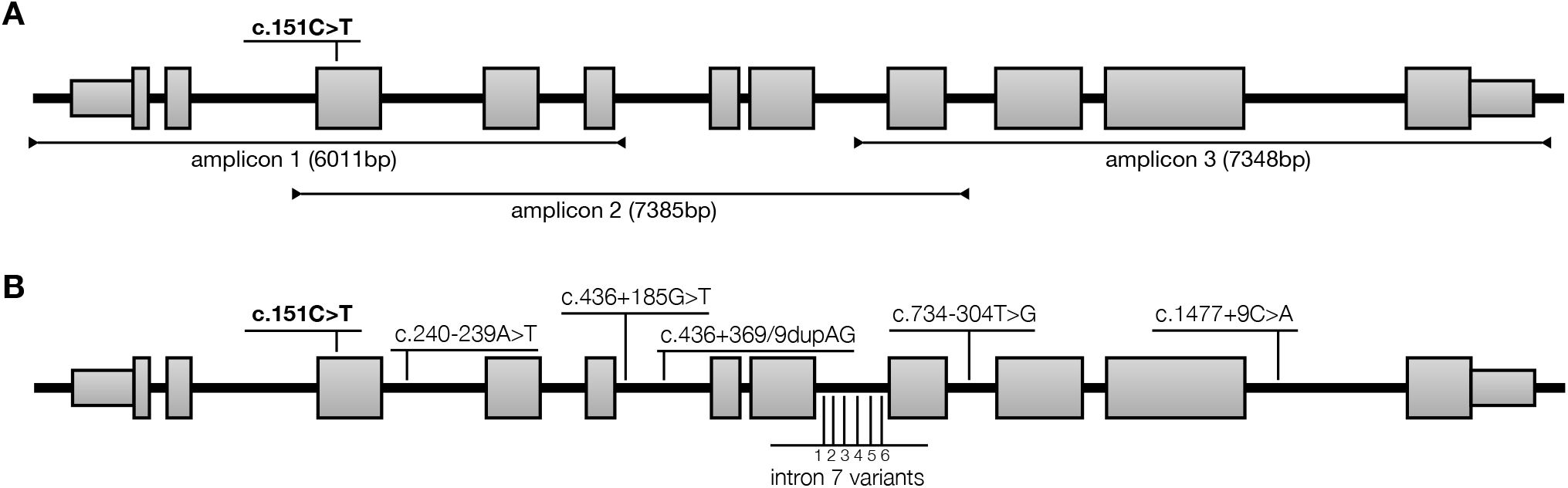
*COCH* haplotype analysis. **A**) Overview of the amplicons used to determine the haplotype-specific variants on the c.151C>T mutant *COCH* allele. Amplicon length is indicated in base pairs (bp) between brackets. **B**) Variants with a low population frequency (< 10%) on the c.151C>T mutant haplotype. The six identified variants in intron 7 are 1: c.629+1186T>C; 2: c.629+1779delC; 3: c.629+1807delA; 4: c.629+1809A>C; 5: c.629+1812A>T; 6: c.630-208A>C. Intron-exon structure of transcript NM_001135058.1 is depicted. The c.151C>T variant, causative for DFNA9, is indicated in bold.

**Table 1.** Identified low-frequency variants on the c.151C>T *COCH* haplotype.

| Location | SNP Identifier | Nucleotide change (c. HGVS) | Amino acid change | Frequency (percentage)<br>GnomAD<br>European non-finnish |
| --- | --- | --- | --- | --- |
| e4 | rs28938175 | c.151C>T | Pro51Ser | T: 0.0032 |
| i4 | rs143609554 | c.240-239A>T |  | T: 5.4 |
| i6 | rs7140538 | c.436+185G>T |  | T: 5.5 |
| i6 | rs10701465 | c.436+368_436+369dupAG |  | dupAG: 5.5 |
| i8 | rs186627205 | c.629+1186T>C |  | C: 5.4 |
| i8 | rs200080665 | c.629+1779delC |  | delC: 5.4 |
| i8 | rs368638521 | c.629+1807delA |  | delA: 5.9* |
| i8 | rs554238963 | c.629+1809A>C |  | C: 9.9* |
| i8 | rs184635675 | c.629+1812A>T |  | T: 5.4 |
| i8 | rs2295128 | c.630-208A>C |  | C: 5.3 |
| i9 | rs28362773 | c.734-304T>G |  | G: 7.2 |
| i11 | rs17097458 | c.1477+9C>A |  | A: 5.4 |

\* no data in GnomAD, frequency data from dbSNP 153

### Design and in-silico analysis of AONs

We selected the c.151C>T founder mutation and the intronic, mutant allele-specific variant c.436+368_436+369dupAG as targets for AON-based therapy. In contrast with the identified single nucleotide changes or deletions, the c.436+368_436+369dupAG variant is the only multi-nucleotide variant that is specific to the mutant allele. Based on this, we hypothesized that AONs directed against this variant can provide the highest allele-specificity. To design AONs, we combined the criteria that are commonly used to design splice-switching AONs with the previously established notion that RNase H1-dependent AONs require a series of nucleotides with DNA-like properties in their central region (Pallan and Egli, 2008; Aartsma-Rus et al., 2009; Slijkerman et al., 2018). All possible AONs were investigated for thermodynamic properties *in silico*, with particular attention for the difference in binding affinity between the mutant and wildtype *COCH* mRNA. Targeting regions of all AONs used in this study are shown in Figure 2A. Note that the difference in binding affinity between the mutant and wildtype *COCH* mRNA was predicted to be larger for the AONs directed against the dupAG variant (c.436+368_436+369dupAG) as compared to those directed against the single nucleotide substation (c.151C>T) (Table S1). The recognition of RNA/DNA duplexes by RNase H1 relates to the nature of the carbohydrate moiety in the AON backbone (2’-ribose vs. 2’-deoxyribose) ^16^. Therefore, AONs were either comprised completely of phosphorothioate (PS)-linked DNA-bases, or of a central “gap” region of PS-DNA bases flanked by wings of 2’-O-methyl-RNA bases (gapmers). The gapmer design is particularly suitable for clinical application as the nuclease-resistant 2’-modified ribonucleotides provide an increased binding affinity and half-life time ^12,17,18^.

**Figure 2.**
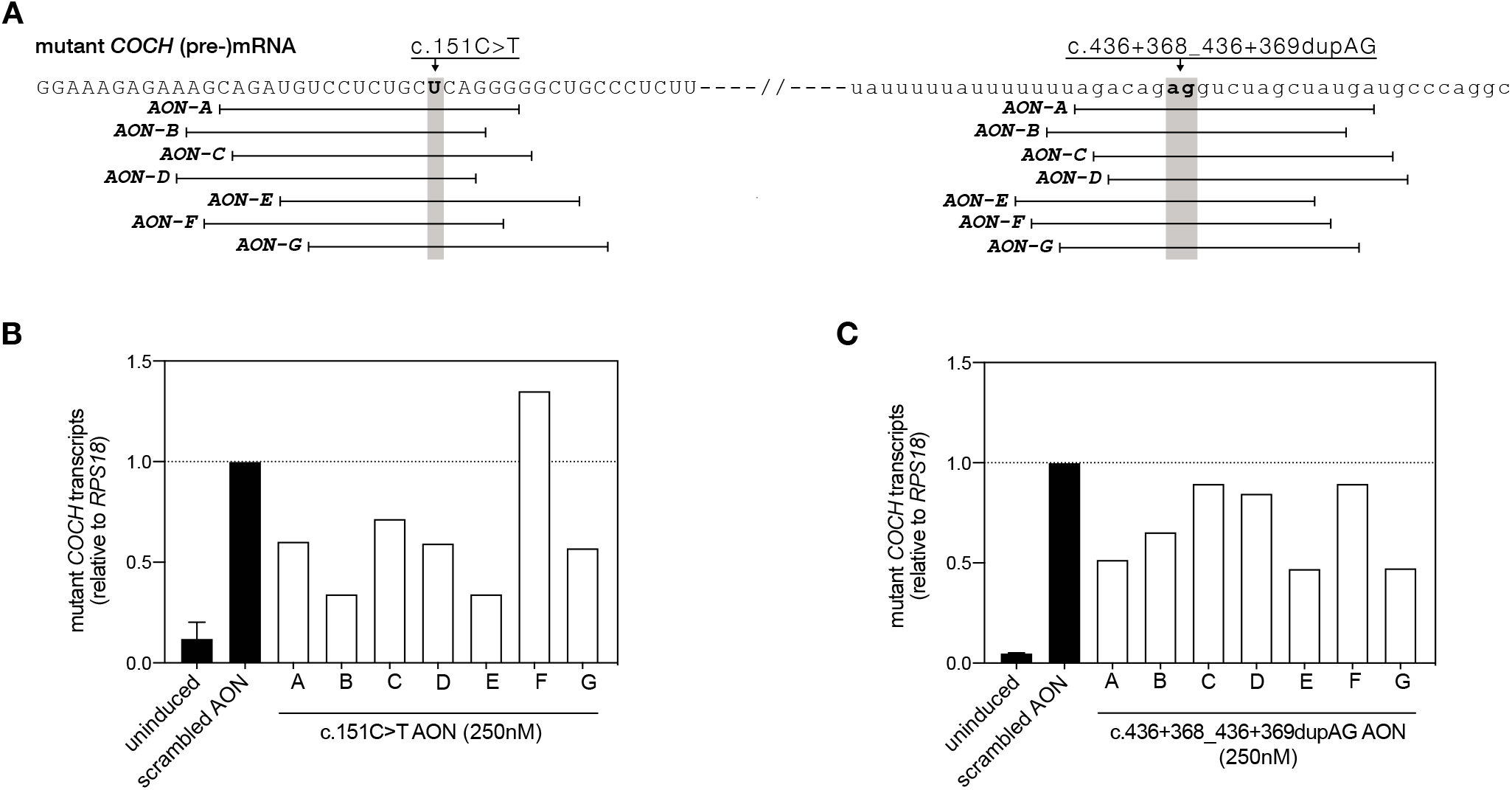
Design and identification of candidate AONs directed against the c.151C>T mutation and the *in cis* intronic variant c.436+368_436+369dupAG. **A**) Graphical representation of AON-RNA binding positions on the c.151C>T mutant *COCH* transcript. Coding sequences are shown in capitals, intronic sequences in lower case. AON sequences are provided in table S1. **B**) Degradation of mutant *COCH* transcripts by AONs (250nM end concentration in the medium), directed against the c.151C>T mutation, in mutant *COCH*-expressing transgenic cells. Six out of the seven AONs were able to lower the levels of mutant *COCH* transcripts at 24 hours post transfection as compared to cells transfected with a scrambled control AON. **C**) Degradation of mutant *COCH* transcripts by different AONs (250nM end concentration in the medium), directed against the c.436+368_436+369dupAG variant on the mutant *COCH* transcript, in mutant *COCH*-expressing transgenic cells. Four out of the seven AONs showed an obvious decrease in mutant *COCH* transcript levels at 24 hours post transfection as compared to cells transfected with a scrambled control AON. Uninduced and scrambled controls are displayed as the average of three biological replicates. Single transfections are used for the screening of on-target AONs. Data are displayed as the fold change compared to scrambled control AON-treated cells, and normalized for the expression of *RPS18.*

### Establishing stable transgenic cell lines expressing wildtype or c.151C>T *COCH* minigenes

The *COCH* expression levels in patient-derived primary fibroblast and Epstein-Barr virus-transformed lymphoblastoid cells are too low to reliably determine the effect of RNase H1-depended antisense oligonucleotides (AONs). Therefore, we used the Flp-In^TM^ system to generate two stable transgenic T-REx^TM^ 293 cell lines, expressing either a mutant (including three deep-intronic allele-discriminating variants; Figure 1) or a wildtype *COCH* minigene construct under the control of a tetracycline-dependent promotor. The minigene constructs span the genomic *COCH* sequence between the transcription initiation site and the last complete nucleotide triplet of exon 7 (Figure S2A). For both alleles, several clones were expanded and investigated for inducible *COCH* expression. For further experiments, wildtype and mutant clones were selected with similar *COCH* expression levels upon activation of the tetracycline-dependent promotor (Figure S2B). Correct pre-mRNA splicing of both wildtype and mutant minigene *COCH* exons 1-7 was confirmed with RT-PCR (Figure S2C). In order to reliably quantify mutant and wildtype *COCH* transcript levels, we used a custom Taqman™ assay (Applied Biosystems) in which different fluorophores are coupled to probes specific for either the mutant or the wildtype transcript.

### RNase H1-dependent antisense oligonucleotides specifically target mutant *COCH* transcripts for degradation

As the *COCH* gene is continuously expressed in the human cochlea, we opted for an experimental design in which *COCH* transcription remains active. To induce *COCH* transcription, seeded cells were treated overnight with tetracycline (0.25μg/ml). Next morning, culture medium was replaced by fresh tetracycline-containing medium, and cells were transfected with the AONs at a final concentration in the medium of 250nM. An initial screening of AONs, revealed that six (out of seven) AONs directed against the c.151C>T mutation (Figure 2B) and four (out of seven) AONs directed against the dupAG variant (Figure 2C) were able to decrease the level of mutant *COCH* transcripts as compared to a scrambled control AON.

Three of the most effective AONs directed against the c.151C>T mutation, and one AON directed against the dupAG variant, were analyzed in more detail using two concentrations of gapmer AONs and multiple technical replications (Figure 3). c.151C>T AON-A was able to induce a significant decrease in mutant *COCH* transcripts at a dose of 250nM (P = 0.02, Tukey’s multiple comparison test), but not at 100nM (Figure 3A). While AON-B showed a stronger effect in comparison to AON-A in the initial screening, the effect sizes of AON-A and –B were very similar in this replication experiment (Figure 3B). A significant decrease of mutant *COCH* transcripts was found at both concentrations (P < 0.0012, Tukey’s multiple comparison test). However, the dose of 250nM AON-B was not able to induce a stronger decrease of mutant *COCH* transcripts as compared to the 100nM dose. The third AON directed against the c.151C>T mutation that was investigated in more detail, AON-E, did show a dose-dependent effect size. At 100nM, the level of mutant *COCH* transcripts was approximately half of the amount of transcripts detected in cells treated with a scrambled control AON (P < 0.0002, Tukey’s multiple comparison test). Mutant *COCH* transcript levels were even further decreased in cells transfected with 250nM of AON-E (P < 0.0001, Tukey’s multiple comparison test). While on average the AONs directed against the dupAG variants appeared slightly less effective in the initial AON screen, transfection of mutant *COCH* minigene expressing cells with dupAG AON-B resulted in a significant decrease in mutant *COCH* transcripts at both concentrations tested (Figure 3D; P < 0.0009, Tukey’s multiple comparison test). The effect size of dupAG AON-B was similar to the effect observed for c.151C>T AON-A and -B.

**Figure 3.**
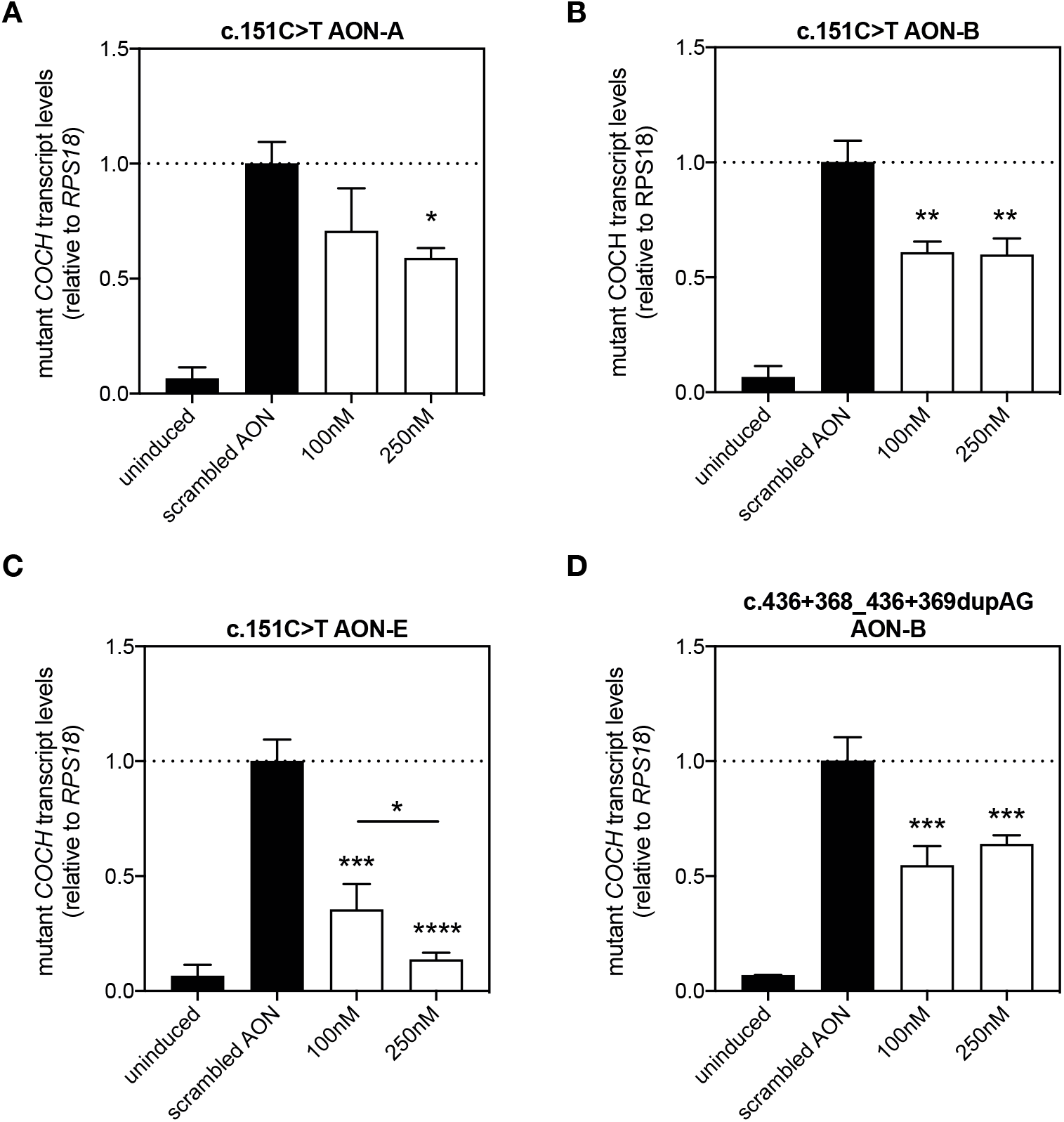
Identified candidate AONs induce a significant decrease in mutant *COCH* transcript levels. To confirm the effect of previously identified candidate AONs c.151C>T AON-A **(A)**, c.151C>T AON-B (**B**), c151C>T AON-E (**C**) and c.436+368_436+369dupAG AON-B (**D**) were investigated at two different doses. **A**) c.151C>T AON-A results in significant decrease in mutant *COCH* transcripts at 250nM, but not at 100nM. **B**) c.151C>T AON-B was able to induce a significant decrease in mutant *COCH* transcripts at both 100nM and 250nM, but no differences between the two doses were observed. **C**) c.151C>T AON-E decreased the level of mutant *COCH* transcripts in a statistically significant and dose-dependent manner. At a concentration of 250nM, the amount of *COCH* transcripts was reduced to 20% of those in cells treated with a scrambled control AON. **D**) Transfection of c.436+368_436+369dupAG AON-B resulted in a significant decrease of mutant *COCH* transcripts, without statistically relevant differences between the two concentrations. All four AONs had a gapmer design with wings of 2’-O-methyl-RNA bases flanking the central PS-DNA core. AONs were transfected at a dose leading to the indicated concentration in the well, and investigated for their effect on transcript levels 24 hours after transfection. Data is expressed as mean ± SD of 3 replicate transfections, normalized to the expression of RPS18 and displayed as the fold change compared to scrambled control AON-treated cells. * P < 0.05, ** P < 0.01, *** P < 0.001, **** P < 0.0001, one-way ANOVA with Tukey’s post-test.

Finally, we investigated the specificity of these four AONs in discriminating between mutant and wildtype *COCH* transcripts (Figure 4). We chose to compare the AONs at a concentration of 100nM, as three out of the four AONs were able to significantly reduce mutant *COCH* transcript levels at this concentration. As observed previously, transfection of mutant *COCH* minigene cells with c.151C>T AON-B, c.151C>T AON-E, and dupAG AON-B, significantly decreased mutant *COCH* transcripts levels as compared to a scrambled control AON (Figure 4A). None of the four AONs induced a significant decrease of wildtype *COCH* transcripts when transfected in wildtype *COCH* expressing transgenic cells, although we did observe a marked decrease in both mutant and wildtype *COCH* transcript levels resulting from the transfection of c.151C>T AON-A (Figure 4B). Likely, the correction for multiple comparisons explains the lack of a significant difference between c.151C>T AON-A treated and scrambled AON treated wildtype *COCH* minigene cells. The results for c.151C>T AON-E are of particular interest, as this gapmer was able to decrease the levels of mutant *COCH* transcripts with almost 60% compared to a scrambled AON, but had no significant effect on the level of wildtype *COCH* transcripts. In addition, dupAG AON-B also displayed perfect allele discrimination, albeit with a smaller effect size on mutant *COCH* transcripts as compared to c.151C>T AON-E.

**Figure 4.**
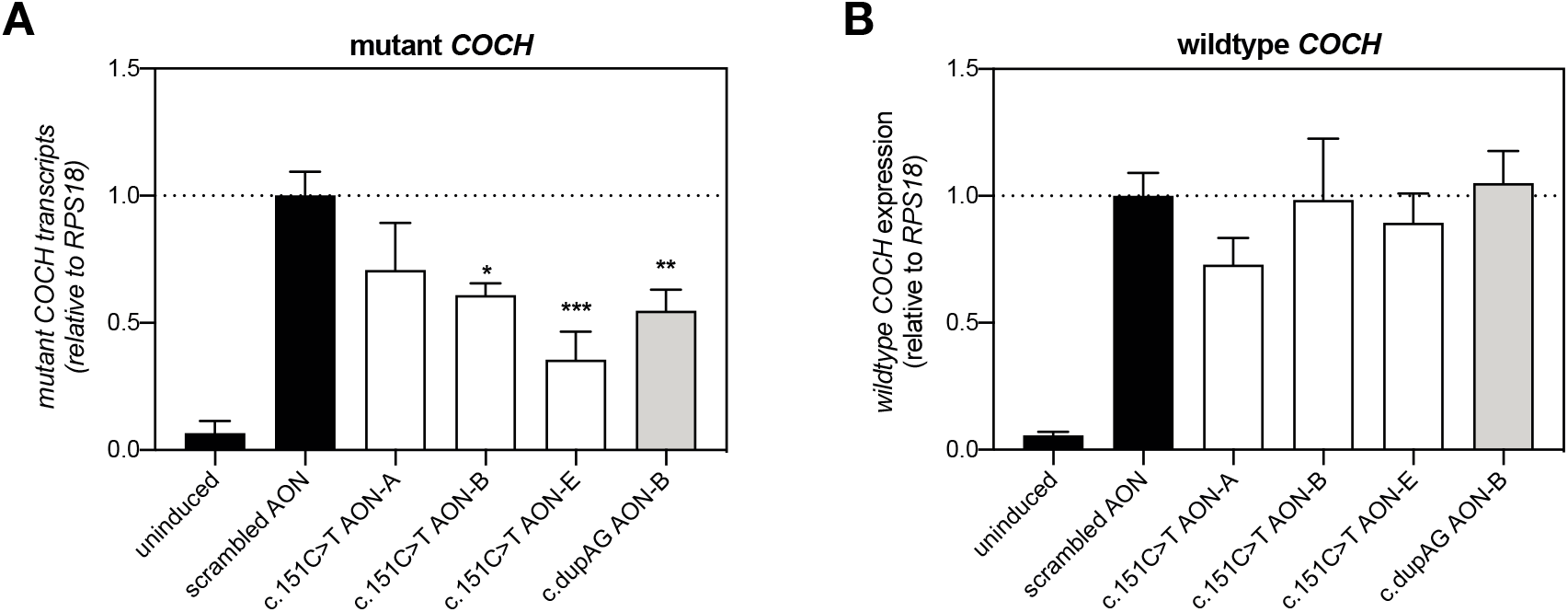
Allele-specificity of the identified AONs. AONs directed against the c.151C>T mutation or the c.436+368_436+369dupAG (dupAG) variant were transfected in stable transgenic cell lines expressing **A**) a mutant *COCH* minigene, and **B**) a wildtype *COCH* minigene. AONs were transfected at a dose that results in a final concentration of 100nM in the culture medium, and their effect on *COCH* transcript levels was investigated 24 hours post transfection. **A**) As shown previously, c.151C>T AON-B and AON-E, and dupAG AON-B, were able to induce a significant decrease in the mutant *COCH* transcript level. **B**) None of the AONs resulted in a significant decrease in wildtype *COCH* transcript levels in transgenic cells expressing the wildtype *COCH* minigene. While c.151C>T AON-A results in a decrease in wildtype *COCH* transcript levels, the observed decrease is not statistically significant (P = 0.14, Tukey’s multiple comparison test). All AONs used here consisted of a gapmer composition. Data are displayed as the fold change compared to untreated cells (mean ± SD) of 3 replicates, and normalized for the expression *RPS18.* * P < 0.05, ** P < 0.01, one-way ANOVA with Tukey’s post-test.

## Discussion

The c.151C>T founder mutation in *COCH* is estimated to be one of most prevalent causes of dominantly-inherited, adult-onset hearing loss and vestibular dysfunction, affecting >1000 individuals in the Dutch/Belgian population. In this work, we present 11 intronic variants *in cis* with the c.151C>T mutation, and show that these variants can be exploited for the development of a mutant allele-specific therapy using RNase H1-dependent antisense oligonucleotides (AONs). We identified a highly effective and mutant-allele specific AON, directed against the c.151C>T mutation, as the most promising candidate for further preclinical development.

The ability of antisense oligonucleotides (AONs) to specifically target mutant transcripts for degradation is of key importance for the development of an AON-based therapy for dominantly-inherited disorders with a dominant-negative or gain-of-function disease mechanism such as DFNA9. The therapeutic strategy must be potent enough to prevent the synthesis of proteins from the mutant allele, but allow sufficient protein synthesis from the wildtype allele for normal inner ear function. For any antisense-based approach, discrimination between alleles based on a single nucleotide difference presents as a potential pitfall in terms of concomitated downregulation of the wildtype allele ^19–21^. Recently published AONs directed against a mutation in *NR2E3,* causative for dominantly inherited retinitis pigmentosa, also significantly reduced the wildtype transcript and protein levels ^22^. In contrast, for Huntington’s disease *(HTT* gene), also resulting from a non-haploinsufficiency disease mechanism, the use of gapmer AONs to target a single nucleotide polymorphism (SNP) specific to the mutant allele emerged as a promising therapeutic strategy *in vitro* and *in vivo* ^15^. Haplotype mapping of candidate SNPs in the *HTT* gene was previously done manually via genotyping of family trios ^23^. As nearly all cases of DFNA9 are caused by single nucleotide changes (Bae et al., 2014), we explored the presence of mutant allele-specific variants that can serve as additional targets to develop a therapeutic strategy for the most frequently occurring DFNA9 mutation c.151C>T. Here, we employed SMRT sequencing ^24^ to sequence the complete mutant *COCH* haplotype using three overlapping PCR amplicons. With average polymerase read lengths of up to 30kb, the SMRT sequencing platform presents a powerful tool to identify genetic variants on the mutant allele.

The c.151C>T *COCH* allele contains a remarkably high number of SNPs with a relatively low allele frequency (~5%) in the non-Finnish European population according to the gnomAD database (v.2.1.1) ^25^. As the c.151C>T founder mutation arose on a relatively uncommon haplotype, we estimate that less than 5% of DFNA9 patients are homozygous for these SNPs. Therefore, approximately 95% of DFNA9 patients with the c.151C>T mutation can be treated with AONs directed against one of these mutant allele-specific variants. In comparison, it was reported that targeting one of three relatively frequent SNPs can provide a treatment for approximately 85% of patients suffering from Huntington’s disease ^23^. In contrast to the identified mutant allele-specific SNPs in *HTT,* all of the identified variants in *COCH* map to the introns. As such, the identified mutant allele-specific variants in *COCH* are only amenable to AON-mediated pre-mRNA degradation by the RNase H1 enzyme, and not to mRNA interference (RNAi) ^26–28^.

We designed AONs to specifically target mutant *COCH* transcripts for RNase H1 degradation. In addition to targeting the DFNA9-associated mutation c.151C>T, we opted to target the 2bp duplication c.436+368_436+369dupAG *in cis* with the DFNA9 mutation. In-silico analysis of thermodynamic AON properties indicated that AONs directed against the dupAG variant possess a larger difference in binding affinity between the mutant and the wildtype transcript as compared to AONs directed against the c.151C>T mutation (Table S1). The on-target and off-target efficacy of AONs was investigated in stable transgenic cells that express mutant or wildtype *COCH* minigenes under control of a tetracycline-dependent promotor. A similar cell model was previously used to investigate the kinetics of RNase H1-dependent antisense oligonucleotide induced degradation ^13^, and offers a suitable alternative to the patient-specific fibroblast and lymphoblastoid cell lines that hardly express *COCH.* We opted to investigate the effect of AONs under continuous activation of *COCH* transcription, which best resembles the situation in the cochlea, where constant *COCH* expression amounts to synthesis of one of the most abundant proteins in the entire organ ^1,6^. The gapmer configuration of c.151C>T AON-E was the most effective of all the designed AON molecules, and at the highest dose resulted in a decrease of mutant *COCH* transcripts to < 15% of the number of transcripts in cells treated with a scrambled control AON. The effect of AONs directed against the c.436+368_436+369dupAG (dupAG) variant was overall lower as compared to the c.151C>T AONs in the initial screening experiment. The effect of dupAG AON-B was also less potent as compared to c.151C>T AON-E. This could result from small differences in biochemical properties. The predicted on-target binding affinity of all AONs directed against the dupAG variant was indeed lower as compared to AONs directed against the c.151C>T mutation. Biochemical properties of the dupAG AONs can be improved by increasing the length of the AON, or by introducing chemically modified nucleotides that enhance binding affinity. However, the lower effect of the dupAG AONs on mutant *COCH* transcript levels could also be related to the fact that these AONs are directed against an intronic variant, which is only present in unspliced nuclear pre-mRNA. In contrast, AONs directed against exonic targets act on all transcripts, both in the nucleus and cytoplasm. With the observed efficiency and high allele-specificity of c.151C>T AON-E, for which therapeutic application is also not constrained by a small percentage of individuals that is homozygous for the target variant, we concluded that there is currently little need to optimize the AONs that target intronic variants.

The transient effect of AONs is both an advantage and a potential limitation for future clinical applications. It lowers the risk of sustained adverse or off-target effects that could accompany genome editing techniques, but it also implies that a repeated delivery is likely to be required to achieve maximum efficacy. AON-based splice-modulation therapy for hearing impairment in Usher syndrome type 1C is extensively investigated in the fetal and post-natal cochlea ^29,30^.

Delprat et al previously reported the use of phosphorothioate oligonucleotide-mediated knockdown to investigate the role of the otospiralin protein in the inner ear protein ^31^. In this study, they placed pieces of gel foam loaded with AONs on the round window membrane (RWM) of rats, and observed the effects of otospiralin knockdown already two days later ^31^. Otospiralin and cochlin are both expressed by the otic fibrocytes, which indicates that cellular uptake of AONs is unlikely to be a limiting factor for DFNA9 therapy. Although many advancements in cochlear drug delivery have been made since (reviewed by *e.g.* ^32–34^, a huge gap in knowledge remains in terms of safety, stability and biodistribution of gapmer AONs in the (adult) human cochlea. Further investigation into the feasibly of RWM diffusion as a potential delivery method for AON-based therapy in patients is also warranted, as the gapmer composition of AONs may affect diffusion properties, and the thickness of the human RWM and larger size of the cochlea are likely to affect the biodistribution of AONs.

The reported age of onset of auditory and vestibular symptoms in c.151C>T DFNA9 in patients, on average in the 3rd or 4th decade of life ^35^, suggests that the inner ear can cope with the burden of mutant cochlin proteins for several decades before it leads to detectable auditory and vestibular damage and dysfunction. It has also been shown that otic fibrocytes, the main cell type expressing cochlin, display some capacity for self-renewal ^36^. In the most optimal situation, AONs might be able to remove the burden of mutant cochlin proteins to an extent that allows for fibrocyte renewal and thereby possibly improved auditory and vestibular function. Halting the disease progression in an early stage is likely a more realistic outcome, and would already greatly improve the patient’s quality of life. Further pre-clinical studies in animal models are therefore not only required to determine both the therapeutic efficacy and allele-specificity on the long term, but also the need and frequency for repeated delivery.

In conclusion, this study shows that AONs can be engineered to specifically target the c.151C>T mutant *COCH* transcript for degradation. Targeting of intronic, mutant-allele-specific variants present an interesting opportunity to further improve efficacy and allele-specificity of AON-based therapy for DFNA9. Models for the long-term investigation of the effects of AONs are not (yet) available. Efficacy studies in appropriate animal models will provide important insights into the feasibility and specificity of AON-based therapy for DFNA9. Combined with the rapidly evolving procedures for repeated drug delivery to the cochlea, the AONs developed in this study form the first step towards the development of a genetic therapy for DFNA9.

## Materials and methods

### Single-Molecule Real-Time (SMRT) sequencing of COCH haplotypes

This study was approved by the medical ethics committee of the Radboud University Medical Center in Nijmegen, the Netherlands and was carried out according to the Declaration of Helsinki. Written informed consent was obtained from all participants. DNA samples of three seemingly unrelated DFNA9 patients carrying the c.151C>T mutation in *COCH* were selected for Single-Molecule Real-Time (SMRT) sequencing (Pacific Biosciences, Menlo Park, CA, USA) to identify shared variants on the mutant allele. The *COCH* gene was amplified in three overlapping amplicons (Figure 1), in which known haplotype-specific variants were anticipated to be present to aid assembly. Fragments were amplified with primers 5’-GAAGTTCGGTTCTCAGGCC-3’ and 5’-TGCCATCGTCATACAAAAGG-3’ (fragment 1), 5’-CAAAATCTGGAATGGTATGGAAG-3’ and 5’-GATCAAATGCAGACCTAGCC-3’ (fragment 2) and 5’-TCCCCTGCAGTACTTTTTGTC-3’ and 5’-GTAAGCCAGCTTACAATAACTC-3’ (fragment 3), using Q5 polymerase (New England Biolabs, Ipswich, MA, USA) according to manufacturer’s instructions. Amplicons were pooled per sample, and library preparation was done according to protocol ‘Procedure and Checklist – Preparing SMRTbell Libraries using PacBio Barcoded Adapters for Multiplex SMRT Sequencing’ (Pacific Biosciences, Part Number 100-538-700-02). Generation of polymerase bound SMRTbell complexes was performed using the Sample Setup option in SMRTLink 6.0 (Pacific Biosciences) and sequencing was performed on a Sequel I systems (Pacific Biosciences). Following the run, generation of circular consensus reads (CCS) and mapping of these reads was performed using SMRTLink 6.0. Bam files were loaded into the Sequence Pilot software (JSI medical systems) to perform variant calling. The variants were subsequently filtered to excluded homopolymers, homozygous variants. The identified variants with a low population frequency (< 10%) were considered as potential therapeutic targets, and validated using targeted sanger sequencing. Segregation analysis in two branches of large Dutch DNFA9 families (W02-006 and W00-330) was used to confirm the presence of the identified variants on the mutant haplotype. Primers used in the segregation analyses are listed in table S2

### Antisense oligonucleotides

Antisense oligonucleotides (AONs) were designed using previously published criteria for splice-modulating AONs ^37,38^. In summary, the sequences surrounding the c.151C>T and c.436+368_436+369dupAG variants on the mutant *COCH* allele were analyzed *in silico* for AON-accessibility. The thermodynamic properties of every possible 20-mer antisense oligonucleotide were analyzed *in silico* for AON-AON duplex formation, the formation of AON-target mRNA duplexes and the formation of AON-wildtype mRNA duplexes using the RNAstructure webserver ^39^. The uniqueness of the AON target sequences was determined by BLAST analysis. The seven most optimal AONs were purchased from Eurogentec (Liège, Belgium) and dissolved in phosphate-buffered saline (PBS) before use. As a non-binding control, an AON with a scrambled nucleotide sequence was also acquired. Sequences and AON chemistry are presented in table S1.

### Generation of transgenic COCH minigene cell lines

The genomic region of wildtype and c.151C>T mutant *COCH* exons 1 to 7 (transcript variant 1; NM_001135058.1), including the haplotype-specific variants, was amplified from the translation initiation site to the splice donor site of exon 7 using primers 5’-ATGTCCGCAGCCTGGATC-3’ and 5’-GGCTTGAACAAGGCCCACA-3’. The mutant and wildtype amplicons were subsequently cloned into the pgLAP1 vector (Addgene plasmid #19702) using Gateway cloning technology (Invitrogen, Carlsbad, USA). Upon sequence validation, *COCH*-containing pgLAP1 vectors were co-transfected with pOGG44 (# V600520, Invitrogen), encoding Flp-Recombinase, in FLp-in^TM^ T-REx^TM^ 293 cells (# R78007, Invitrogen) using polyethylenimine. Cells in which the *COCH* sequence was stably integrated were selected for using DMEM-AQ medium (Sigma Aldrich, Saint Louis, USA) supplemented with 10% Fetal Calf Serum, 1% Penicillin/Streptomycin, Sodium Pyruvate, 10ug/ml blasticidin and 100ug/ml hygromycin. Individual hygromycin-resistant clones were expanded and subsequently tested for the induction of *COCH* transcription by tetracycline using an allele-specific TaqMan™ assay. Correct splicing of the *COCH* minigenes was assessed using a forward primer on exon 1 (5’-TCCGCAGCCTGGATCCCGG-3’) and reverse primer on exon 7 (5’-GGCTTGAACAAGGCCCACA-3’).

### Delivery of RNase H1-dependent antisense oligonucleotides

Wildtype and mutant *COCH*-expressing FLp-in^TM^ T-REx^TM^ 293 cells were cultured in DMEM-AQ medium (Sigma Aldrich, Saint Louis, USA) supplemented with 10% Fetal Calf Serum, 1% Penicillin/Streptomycin, Sodium Pyruvate, 10ug/ml blasticidin and 100ug/ml hygromycin. For AON treatments, cells were seeded in 12-well or 24-well plates at ~50% confluency. Next day, *COCH* transcription was activated through the administration of 0.25 μg/ml tetracycline (# T7660, Sigma Aldrich). Twenty hours after induction, cells were transfected with AONs using Lipofectamine 2000 (Invitrogen) according to manufacturer’s instructions, using a 1:2 ratio of AON (in μg) and lipofectamine reagent (in μl). AON doses are calculated as final concentration in the culture medium. Cells were collected for transcript analysis 24 hours after AON delivery.

### RNA extraction and cDNA synthesis

Total RNA was extracted from cells using the Nucleospin RNA mini kit (# 740955, Machery-Nagel) according to manufacturer’s instructions. First strand cDNA was generated using iScript cDNA synthesis reagents (Bio-Rad, Hercules, USA) using a fixed amount of RNA input (250ng) in a 10ul reaction volume. The obtained cDNA was diluted four times and used for transcript analysis.

### Analysis of COCH transcript levels

Diluted cDNA (4μl) was used as input in an allele-specific TaqMan assay using primers 5’-GGACATCAGGAAAGAGAAAGCAGAT-3’ and 5’-CCCATACACAGAGAATTCCTCAAGAG-3’, a wildtype allele-specific VIC-labeled probe 5’-CCCCCTGGGCAGAG-3’ and a mutant allele-specific FAM-labeled probe 5’-CCCCCTGAGCAGAG-3’. Expression of *RPS18* was analyzed with GoTaq (# A6002, Promega), using primers 5’-ATACAGCCAGGTCCTAGCCA-3’ and 5’-AAGTGACGCAGCCCTCTATG-3’. Abundance of mutant and wildtype *COCH* transcripts was calculated relative to the expression of the housekeeping gene *RPS18.*

## Acknowledgments

This work is financially supported by the Dutch Organization for Scientific research (NWO Offroad grant 40-08125-98-16065; E.d.V) and the Queen Elisabeth Medical Foundation for Neurosciences (E.d.V and E.v.W.). Patent has been filed for the AONs described in this manuscript under number 19206490.5. E.d.V. and E.v.W. report being employed by Radboudumc and inventor on this patent. SMRT sequencing was done at the Radboudumc Genome Technology Center.

## Author Contributions

Conceptualization: E.d.V. and E.v.W.; Methodology: E.d.V. and J.P.; Formal analysis: E.d.V.; Investigation: E.d.V., J.P., J.C.M., A.A.M, J.O., S.v.d.H.; Resources: E.d.V, K.N., R.P. and E.v.W.; Writing – original draft: E.d.V.; Writing – review and editing: H.K. and E.v.W., Supervision: E.d.V, E.vW and H.K.

---

**Figure S1.**
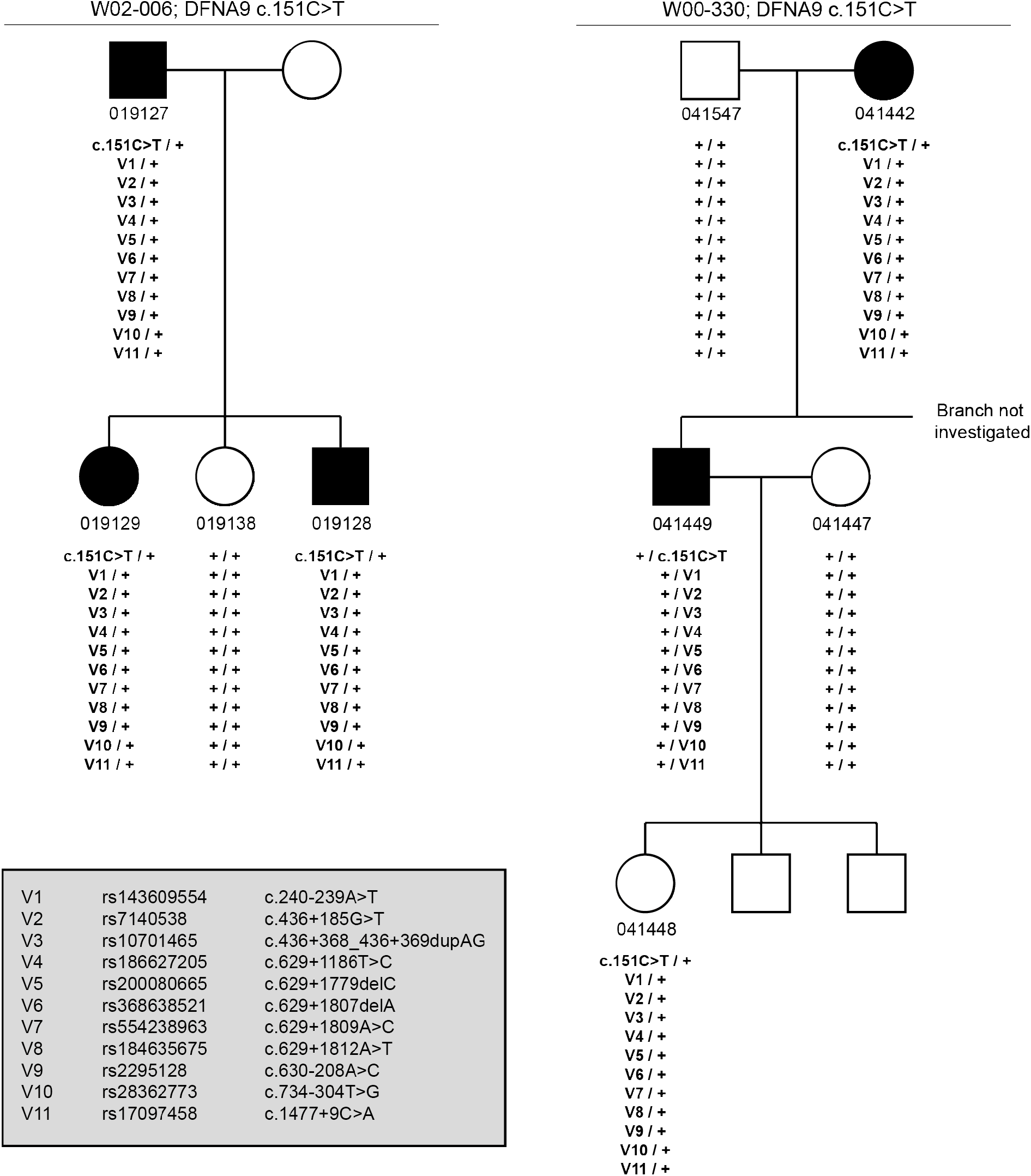
Segregation analysis of haplotype-specific variants. Small branches from the pedigrees of two large Dutch DFNA9 families (W02-006 and W00-330) were investigated to confirm co-segregation of the haplotype-specific variants with the c.151C>T mutation. Numbers below each individual depict the internal identifier of the DNA samples. Individual 041448 was not clinically affected at the time of sample collection. V1-V10: *COCH* variants (see grey box); +: wildtype; square: male; circle: female; open symbol: clinically unaffected; closed symbol: clinically affected.

**Figure S2.**
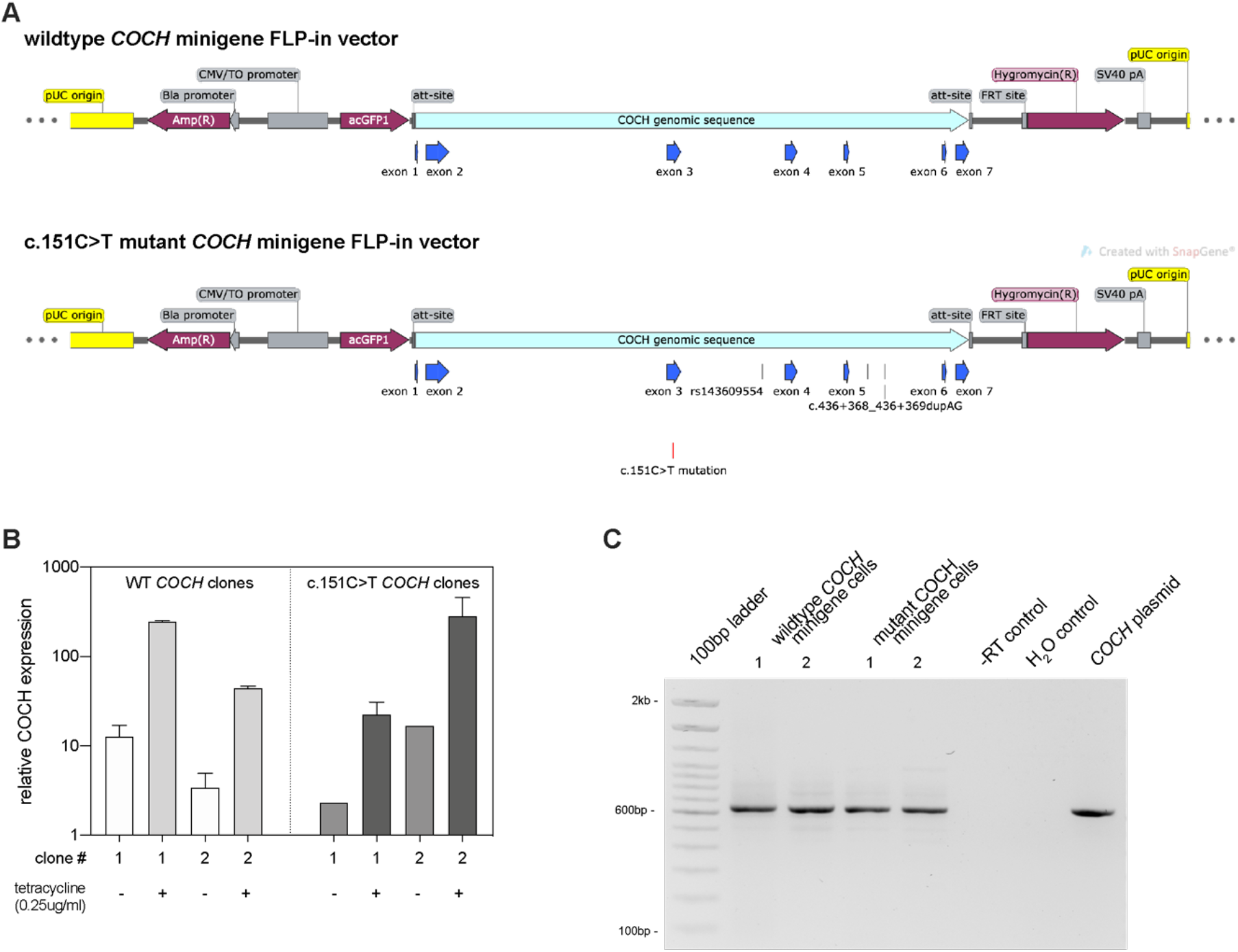
Inducible *COCH* minigene T-REx 293T cells. **A)** schematic overview of the wildtype and mutant COCH vectors that were used to establish the COCH minigene T-REx 293T cells. **B)** Measurement of *COCH* expression upon overnight induction with tetracycline. Two clones of wildtype *COCH* minigene-expressing transgenic cells, and two clones of mutant *COCH* minigene-expressing transgenic cells were investigated. Wildtype clone 2, and mutant clone 1 were selected for experiments based on the relatively similar levels of *COCH* expression upon tetracycline treatment. Note that uninduced cells always show a certain level of background *COCH* expression. As the Taqman™ probe for the mutant *COCH* transcript is highly specific, it appears that the transcriptional activity of the tetracycline promotor is not completely off in uninduced cells. Data shown as mean ± SD. **C)** RT-PCR analysis of *COCH* transcripts in tetracycline-treated mutant and wildtype *COCH* minigene-expressing cells. For each cell line, two replicate samples are shown. Sanger sequencing of the amplicons confirmed correct splicing of the minigene *COCH* transcripts. The positive control is a plasmid containing the coding sequence of *COCH* that was amplified from fetal cochlear cDNA.

## Reference list

1. Robertson, NG, Cremers, CWRJ, Huygen, PLM, Ikezono, T, Krastins, B, Kremer, H, et al. (2006). Cochlin immunostaining of inner ear pathologic deposits and proteomic analysis in DFNA9 deafness and vestibular dysfunction. Hum Mol Genet 15: 1071–1085.

2. Bom, SJH, Kemperman, MH, Huygen, PLM, Luijendijk, MWJ and Cremers, CWRJ (2003). Cross-sectional analysis of hearing threshold in relation to age in a large family with cochleovestibular impairment thoroughly genotyped for DFNA9/COCH. Ann. Otol. Rhinol. Laryngol. 112: 280–286.

3. De Belder, J, Matthysen, S, Claes, AJ, Mertens, G, Van de Heyning, P and Van Rompaey, V (2017). Does Otovestibular Loss in the Autosomal Dominant Disorder DFNA9 Have an Impact of on Cognition? A Systematic Review. Front Neurosci 11: 735.

4. Gallant, E, Francey, L, Fetting, H, Kaur, M, Hakonarson, H, Clark, D, et al. (2013). Novel COCH mutation in a family with autosomal dominant late onset sensorineural hearing impairment and tinnitus. Am J Otolaryngol 34: 230–235.

5. Jung, J, Yoo, JE, Choe, YH, Park, SC, Lee, HJ, Lee, HJ, et al. (2019). Cleaved Cochlin Sequesters Pseudomonas aeruginosa and Activates Innate Immunity in the Inner Ear. Cell Host & Microbe 25: 513–525.e6.

6. Nagy, I, Trexler, M and Patthy, L (2008). The second von Willebrand type A domain of cochlin has high affinity for type I, type II and type IV collagens. Febs Lett 582: 4003–4007.

7. de Kok, YJ, Bom, SJ, Brunt, TM, Kemperman, MH, van Beusekom, E, van der Velde-Visser, SD, et al. (1999). A Pro51Ser mutation in the COCH gene is associated with late onset autosomal dominant progressive sensorineural hearing loss with vestibular defects. Hum Mol Genet 8: 361–366.

8. Fransen, E, Verstreken, M, Bom, SJ, Lemaire, F, Kemperman, MH, de Kok, YJ, et al. (2001). A common ancestor for COCH related cochleovestibular (DFNA9) patients in Belgium and The Netherlands bearing the P51S mutation. J. Med. Genet. 38: 61–65.

9. Yao, J, Py, BF, Zhu, H, Bao, J and Yuan, J (2010). Role of protein misfolding in DFNA9 hearing loss. Journal of Biological Chemistry 285: 14909–14919.

10. Jones, SM, Robertson, NG, Given, S, Giersch, ABS, Liberman, MC and Morton, CC (2011). Hearing and vestibular deficits in the Coch(-/-) null mouse model: comparison to the Coch(G88E/G88E) mouse and to DFNA9 hearing and balance disorder. Hear. Res. 272: 42–48.

11. JanssensdeVarebeke, SPF, Van Camp, G, Peeters, N, Elinck, E, Widdershoven, J, Cox, T, et al. (2018). Bi-allelic inactivating variants in the COCH gene cause autosomal recessive prelingual hearing impairment. Eur. J. Hum. Genet. 23: 42.

12. Crooke, ST (1999). Molecular mechanisms of action of antisense drugs. Biochimica et Biophysica Acta (BBA) - Gene Structure and Expression 1489: 31–43.

13. Vickers, TA and Crooke, ST (2015). The rates of the major steps in the molecular mechanism of RNase H1-dependent antisense oligonucleotide induced degradation of RNA. Nucleic Acids Res 43: 8955–8963.

14. Vickers, TA and Crooke, ST (2014). Antisense Oligonucleotides Capable of Promoting Specific Target mRNA Reduction via Competing RNase H1-Dependent and Independent Mechanisms. PLoS One 9: e108625.

15. Southwell, AL, Kordasiewicz, HB, Langbehn, D, Skotte, NH, Parsons, MP, Villanueva, EB, et al. (2018). Huntingtin suppression restores cognitive function in a mouse model of Huntington’s disease. Sci Transl Med 10: eaar3959.

16. Pallan, PS and Egli, M (2008). Insights into RNA/DNA hybrid recognition and processing by RNase H from the crystal structure of a non-specific enzyme-dsDNA complex. Cell Cycle 7: 2562–2569.

17. Lima, WF and Crooke, ST (1997). Binding affinity and specificity of Escherichia coli RNase H1: impact on the kinetics of catalysis of antisense oligonucleotide-RNA hybrids. Biochemistry 36: 390–398.

18. Bennett, CF and Swayze, EE (2010). RNA targeting therapeutics: molecular mechanisms of antisense oligonucleotides as a therapeutic platform. Annu. Rev. Pharmacol. Toxicol. 50: 259–293.

19. Zaleta-Rivera, K, Dainis, A, Ribeiro, AJS, Cordero, P, Rubio, G, Shang, C, et al. (2019). Allele-Specific Silencing Ameliorates Restrictive Cardiomyopathy Attributable to a Human Myosin Regulatory Light Chain Mutation. Circulation 140: 765–778.

20. Jiang, J, Wakimoto, H, Seidman, JG and Seidman, CE (2013). Allele-specific silencing of mutant Myh6 transcripts in mice suppresses hypertrophic cardiomyopathy. Science 342: 111–114.

21. Southwell, AL, Skotte, NH, Kordasiewicz, HB, Østergaard, ME, Watt, AT, Carroll, JB, et al. (2014). In vivo evaluation of candidate allele-specific mutant huntingtin gene silencing antisense oligonucleotides. Mol. Ther. 22: 2093–2106.

22. Naessens, S, Ruysschaert, L, Lefever, S, Coppieters, F and De Baere, E (2019). Antisense Oligonucleotide-Based Downregulation of the G56R Pathogenic Variant Causing NR2E3-Associated Autosomal Dominant Retinitis Pigmentosa 10: 363.

23. Carroll, JB, Warby, SC, Southwell, AL, Doty, CN, Greenlee, S, Skotte, N, et al. (2011). Potent and selective antisense oligonucleotides targeting single-nucleotide polymorphisms in the Huntington disease gene / allele-specific silencing of mutant huntingtin. Mol. Ther. 19: 2178–2185.

24. Eid, J, Fehr, A, Gray, J, Luong, K, Lyle, J, Otto, G, et al. (2009). Real-time DNA sequencing from single polymerase molecules. Science 323: 133–138.

25. Karczewski, KJ, Francioli, LC, Tiao, G, Cummings, BB, Alföldi, J, Wang, Q, et al. (2019). Variation across 141,456 human exomes and genomes reveals the spectrum of loss-of-function intolerance across human protein-coding genes. bioRxiv 42: 531210.

26. Vickers, TA, Koo, S, Bennett, CF, Crooke, ST, Dean, NM and Baker, BF (2003). Efficient Reduction of Target RNAs by Small Interfering RNA and RNase H-dependent Antisense Agents. J Biol Chem 278: 7108–7118.

27. Østergaard, ME, Kumar, P, Nichols, J, Watt, A, Sharma, PK, Nielsen, P, et al. (2015). Allele-Selective Inhibition of Mutant Huntingtin with 2-Thio-and C5-Triazolylphenyl-Deoxythymidine-Modified Antisense Oligonucleotides. Nucleic Acid Ther 25: 266–274.

28. Smith, CIE and Zain, R (2019). Therapeutic Oligonucleotides: State of the Art. Annu. Rev. Pharmacol. Toxicol. 59: 605–630.

29. Lentz, JJ, Jodelka, FM, Hinrich, AJ, McCaffrey, KE, Farris, HE, Spalitta, MJ, et al. (2013). Rescue of hearing and vestibular function by antisense oligonucleotides in a mouse model of human deafness. Nat Med 19: 345–350.

30. Hastings, ML and Brigande, JV (2020). Fetal gene therapy and pharmacotherapy to treat congenital hearing loss and vestibular dysfunction. Hear. Res.: 107931 doi:10.1016/j.heares.2020.107931.

31. Delprat, B, Boulanger, A, Wang, J, Beaudoin, V, Guitton, MJ, Ventéo, S, et al. (2002). Downregulation of otospiralin, a novel inner ear protein, causes hair cell degeneration and deafness. J. Neurosci. 22: 1718–1725.

32. Plontke, SK and Salt, AN (2018). Local drug delivery to the inner ear: Principles, practice, and future challenges. Hear. Res. 368: 1–2.

33. Chin, OY and Diaz, RC (2019). State-of-the-art methods in clinical intracochlear drug delivery. Curr Opin Otolaryngol Head Neck Surg 27: 381–386.

34. Hao, J and Li, SK (2019). Inner ear drug delivery: Recent advances, challenges, and perspective. Eur J Pharm Sci 126: 82–92.

35. JanssensdeVarebeke, S, Topsakal, V, Van Camp, G and Van Rompaey, V (2019). A systematic review of hearing and vestibular function in carriers of the Pro51Ser mutation in the COCH gene. Eur Arch Otorhinolaryngol 17: 751–12.

36. Bohnenpoll, T, Trowe, M-O, Wojahn, I, Taketo, MM, Petry, M and Kispert, A (2014). Canonical Wnt signaling regulates the proliferative expansion and differentiation of fibrocytes in the murine inner ear. Dev. Biol. 391: 54–65.

37. Aartsma-Rus, A, van Vliet, L, Hirschi, M, Janson, AAM, Heemskerk, H, de Winter, CL, et al. (2009). Guidelines for antisense oligonucleotide design and insight into splice-modulating mechanisms. Mol. Ther. 17: 548–553.

38. Slijkerman, R, Kremer, H and van Wijk, E (2018). Antisense Oligonucleotide Design and Evaluation of Splice-Modulating Properties Using Cell-Based Assays. Methods Mol. Biol. 1828: 519–530.

39. Reuter, JS and Mathews, DH (2010). RNAstructure: software for RNA secondary structure prediction and analysis. BMC Bioinformatics 11: 1–9.

